# An Epilepsy-Causing Mutation Leads to Co-Translational Misfolding

**DOI:** 10.1101/2020.11.25.398222

**Authors:** Janire Urrutia, Alejandra Aguado, Carolina Gomis-Perez, Arantza Muguruza-Montero, Oscar R. Ballesteros, Jiaren Zhang, Eider Nuñez, Covadonga Malo, Hee Jung Chung, Aritz Leonardo, Aitor Bergara, Alvaro Villarroel

## Abstract

Protein folding to the native state is particularly relevant in human diseases where inherited mutations lead to structural instability, aggregation and degradation. In general, the amino acid sequence carries all the necessary information for the native conformation, but the vectorial nature of translation can determine the folding outcome. Calmodulin (CaM) recognizes the properly folded Calcium Responsive Domain (CRD) of K_v_7.2 channels. Within the IQ motif (helix A), the W344R mutation found in epileptic patients has negligible consequences for the structure of the complex as monitored by multiple *in vitro* binding assays and molecular dynamic computations. *In silico* studies revealed two orientations of the side chain, which are differentially populated by WT and W344R variants. Binding to CaM is impaired when the mutated protein is produced *in cellulo* but not *in vitro*, suggesting that this mutation impedes proper folding during translation within the cell by forcing the nascent chain to follow a folding route that leads to a non-native configuration, and thereby generating non-functional ion channels that fail to traffic to proper neuronal compartments.

## INTRODUCTION

Mutations at the *KCNQ2* gene underlie early-onset hereditary epilepsy, with different clinical outcomes (including Benign Familial Neonatal Epilepsy, BFNE and Epileptic Encephalopathy type 7, EE7) [3–7]. This gene encodes for K_v_7.2 subunits of tetrameric voltage-dependent potassium (K^+^) selective channels, which, combined with K_v_7.3 subunits, underlie non-inactivating M-current. K_v_7.2/K_v_7.3 channels are enriched at the plasma membrane of the axon initial segment (AIS) and distal axons [8]. With their characteristic slow voltage-dependent kinetics of activation and deactivation, and voltage-dependent opening within the subthreshold range of action potential generation, they are critical for neuronal excitability [9]. A number of pathogenic variants cluster at key functional domains that are involved in voltage sensing, ion conduction, selectivity, gating, or stabilization of binding to the essential co-factor PIP_2_ [5; 10; 11].

Clusters of pathological variants are also found at the Calcium Responsive Domain (CRD), located intracellularly following the pore gate [5; 12; 13]. The CRD is an autonomously folding hairpin domain formed by two antiparallel alpha helices [14; 15], named A and B [16], that run under the membrane adjacent to the voltage sensor [17]. These helices are recognized by calmodulin (CaM) [18–21], which confers calcium (Ca^2+^) sensitivity [15; 22–26].

Some mutations in the *KCNQ2* gene have been suggested to cause misfolding [27; 28], a mechanism that has been identified in many hereditary diseases [29; 30]. However, little is known on how pathogenic variants located at the CRD affect folding. In this work, we have explored how a K_v_7.2 mutation (W344R) located in helix A of the CRD, found in patients with hereditary epilepsy [1; 31], interferes with channel function. The data reveal that the key mechanism involves the failure to adopt a configuration that can be recognized by CaM *in vivo* but not *in vitro*.

## RESULTS

### Functional characterization of the W344R mutation

The W344R mutation at the IQ site of helix A does not disturb CaM binding to the autonomously folding calcium responsive domain (CRD) of K_v_7.2 channels, yet it abolishes function [1]. This is in contrast to other helix A mutants, for which a clear correlation between CaM binding and function was observed [2](Supplemental Figure 1). To understand this remarkable discrepancy, we re-evaluated its functional properties. In total agreement with previous work [1; 31], we found that homomeric W344R channels were not functional (Figure 1A). The impact of this mutant in combination with its partner K_v_7.3 and wild type (WT) K_v_7.2 subunits (in a 1:2:1 ratio) to mimic genetic balance has been previously tested, revealing variable effects [1; 31]. To simplify the paradigm, we tested the impact of K_v_7.2 mutant when combined with K_v_7.3 subunits in a 1:1 ratio (Figure 1). K_v_7.3 alone yields negligible currents when expressed in HEK293T cells, whereas robust currents, that are two or three fold larger than from homomeric K_v_7.2 currents, are recorded when combined with WT K_v_7.2 subunits in a 1:1 ratio (not shown).

**Figure 1.**
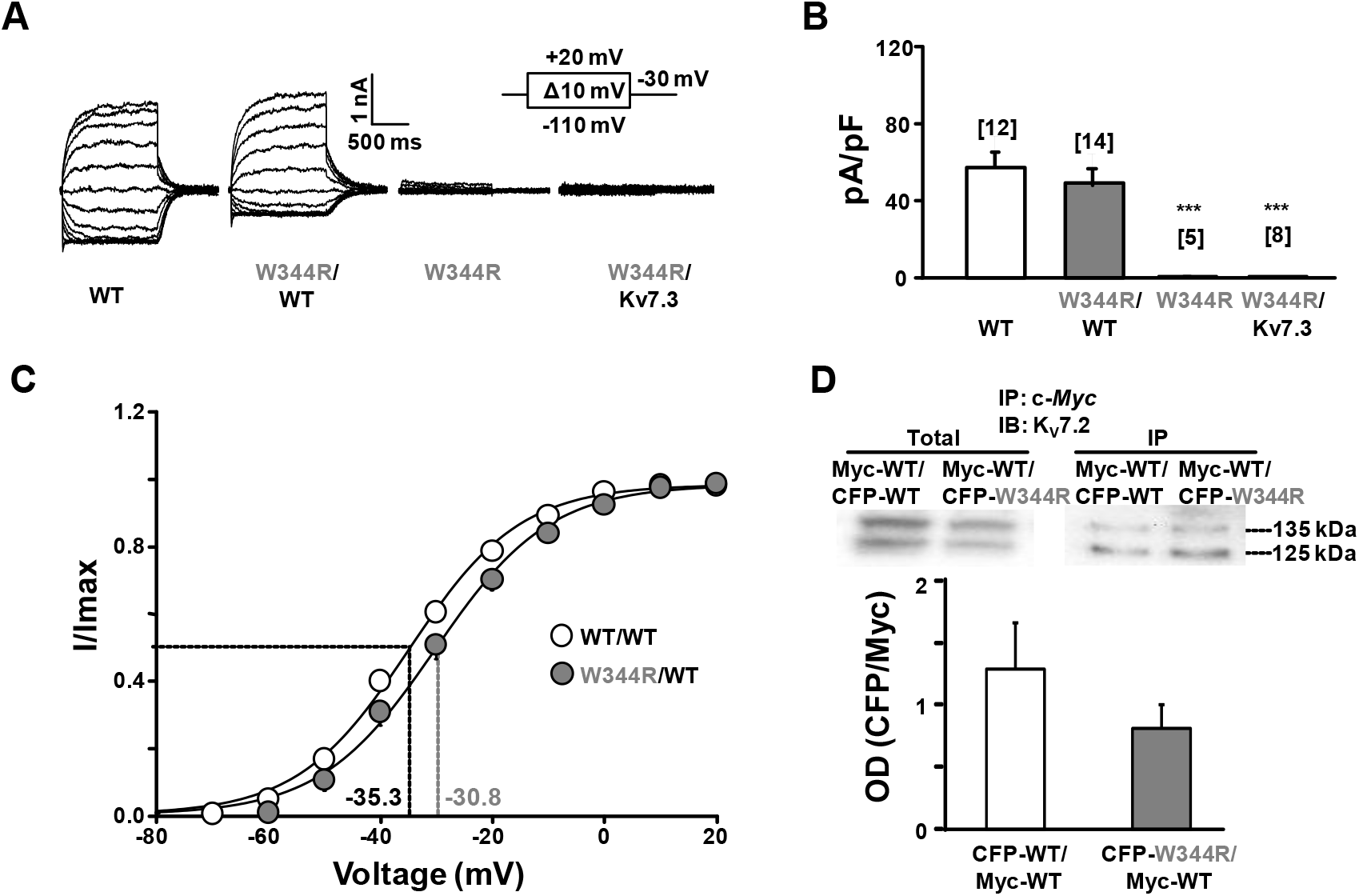
Functional consequences of the W344R mutation. **A.** Representative current traces evoked in cells expressing the indicated subunits. Inset: voltage protocol. **B.** Representation of the current density (pA/pF) from each group at +20 mV from tail currents. K_v_7.2 wild type homomers (WT, white) and heteromers (WT/W344R, gray) had 57.0 ± 11.74 pA/pF and 49.0 ± 8.51 pA/pF, respectively. K_v_7.2 mutant (W344R) and K_v_7.2-W344R/K_v_7.3 heteromers produced negligible current. Number of cells is indicated in brackets. **C.** Current-voltage relationship of K_v_7.2 WT homomers (V_1/2_ = −35.3 ± 0.43 mV, k = 10.5; n = 12) and WT/W344R heteromers (V_1/2_ = −30.8 ± 0.64 mV, k = 11.01; n = 14). ***p < 0.001. **D.** Top: representative immunoblot revealed with anti-K_v_7.2 antibody before (left) and after (right) immunoprecipitating with anti-Myc antibody from cells expressing the indicated Myc-tagged K_v_7.2 subunits (~125 kDa) and CFP-tagged K_v_7.2 (~135 kDa). Bottom: relative densitometry of the two bands. IP: immunoprecipitation, IB: immunobloting. n = 3.

No currents were observed in cells expressing K_v_7.3/K_v_7.2-W344R heteromers, but, in contrast, robust currents were evoked for the K_v_7.2/K_v_7.2-W344R combination (Figures 1A and 1B). There was a modification on the voltage-dependency, as revealed by a statistically significant ~5 mV rightward shift in the current-voltage relationship for the K_v_7.2/K_v_7.2-W344R arrangement (Figure 1). Co-IP experiments were also consistent with the formation of heteromeric assemblies (Figure 1D). HEK293T cells were co-transfected with Myc-tagged K_v_7.2 WT subunits with either CFP-tagged WT or CFP-tagged mutant subunits in a 1:1 ratio. Protein lysates were immunoprecipitated with anti-c-Myc antibody and detected using anti-K_v_7.2 antibody, resulting in the appearance of two bands due to the different MW imposed by the tags (Figure 1D). No significant difference could be detected between the relative signal corresponding of Myc-WT (MW ~125 kDa) and CFP-tagged WT or W344R subunits (MW ~135 kDa).

### The W344R mutation impairs trafficking

K_v_7.2 interaction with CaM is critical for the exit of K_v_7.2/K_v_7.3 channels from the endoplasmic reticulum (ER) and their expression at the axonal surface [8]. To assess the impact on trafficking to the axon initial segment (AIS) and other neuronal regions, surface immunostaining of K_v_7.3 subunits containing an extracellular HA epitope was evaluated [13; 32]. This K_v_7.3 reporter subunit was co-expressed with CFP-tagged K_v_7.2 WT or mutant W344R subunits in embryonic rat hippocampal neurons. As reference, the CFP-tagged K_v_7.2-I340E mutant was included because it precludes CaM binding [32; 33] and, as a consequence, prevents the reporter K_v_7.3 subunit to traffic to the plasma membrane of different neuronal sub-compartments [32]. Plasma membrane expression of HA-K_v_7.3/K_v_7.2 WT heteromers were enriched at the AIS and distal axon (Figure 2). However, when K_v_7.2 carried the I340E or W344R mutations, the reporter HA-K_v_7.3 subunit failed to reach the AIS and distal axon (Figure 2). The total signal for both mutant subunits was reduced by about 70% (supplemental Figures 2 and 3). Except for the relative abundance at the soma (supplemental Figures 2 and 3), the surface expression profile for W344R and I340E mutants were similar (Figure 2), suggesting that neither bind CaM in a cellular *in vivo* context.

**Figure 2.**
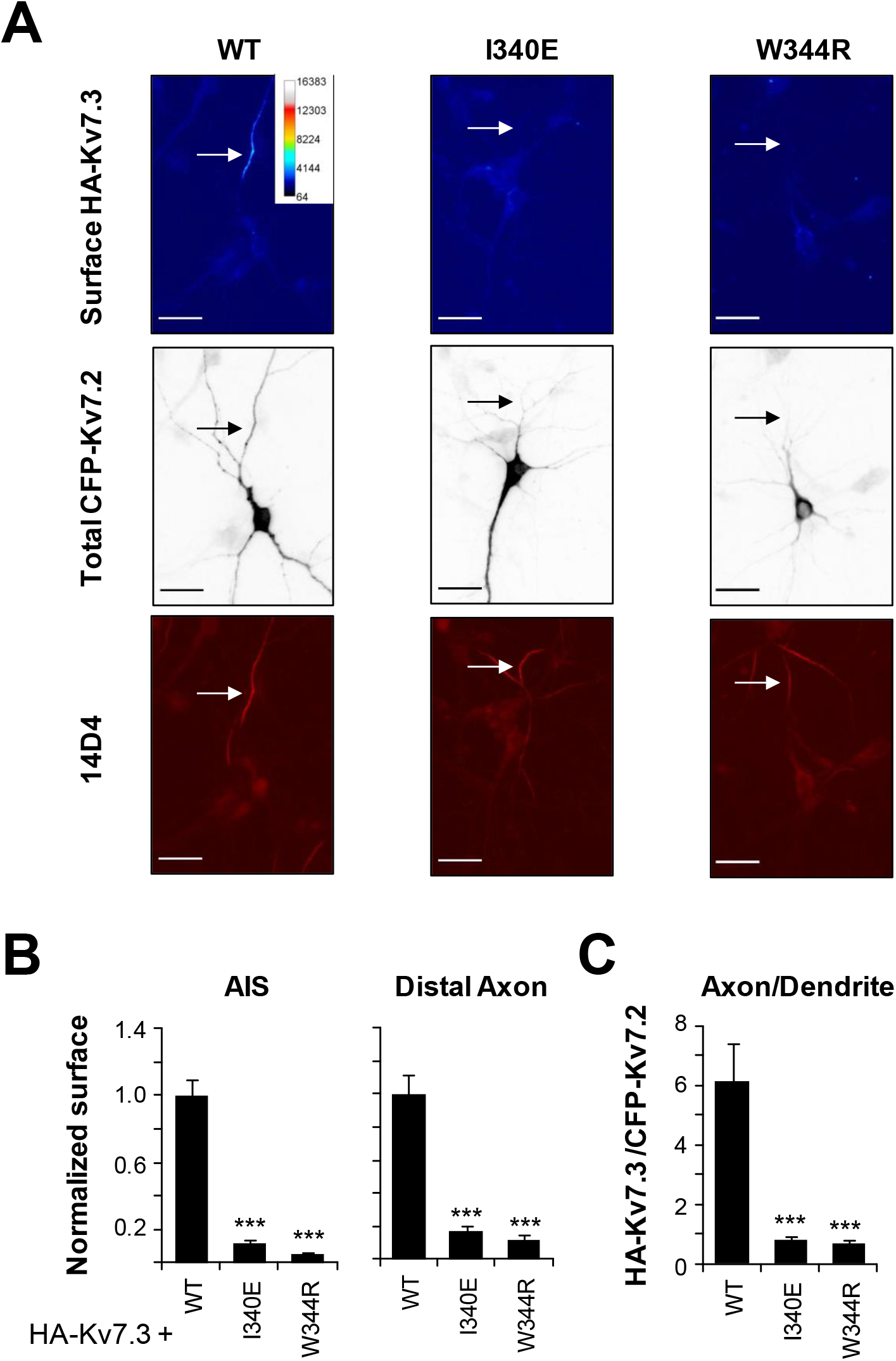
The W344R mutant severely reduced surface expression of heteromeric HA-K_v_7.3/K_v_7.2 in the axons of cultured hippocampal neurons. **A.** Representative images of surface HA-K_v_7.3 as pseudo-color (upper panel), intracellular CFP-K_v_7.2 WT or mutants (middle), and the AIS marker 14D4 (bottom) from neurons transfected with HA-K_v_7.3 and CFP-K_v_7.2 WT, I340E or W344R. Pseudo-color represents different levels of surface HA-K_v_7.3 signal intensity as indicated by the calibration bar. Arrow indicates the locations of AIS. Scale bar: 25 μm. **B.** Background-subtracted fluorescent intensities of surface HA-K_v_7.3 from transfected neurons were normalized to those of HA-K_v_7.3/CFP-K_v_7.2-WT. **C.** The Axon/Dendrite ratio was computed for surface HA-K_v_7.3/CFP-K_v_7.2 fluorescent intensities. Sample numbers for (B-C) are: WT (n = 19), I340E (n = 14), and W344R (n = 13). ***p < 0.005.

### The W344R mutation impairs calmodulin binding *in cellulo*

To assess CaM binding *in cellulo*, the transfer of energy between CaM and K_v_7.2 channels, tagged with CFP (donor) and YFP (acceptor) fluorophores respectively, was monitored in living cells using Förster resonance energy transfer (FRET). This pair of fluorophores exhibits 50% energy transfer at ~50 Å and produces measurable transfer up to ~80 Å. Binding between a ligand and acceptor can be evaluated *in cellulo* using this approach, with comparable results to *in vitro* binding assays [34–36]. Cells expressing similar levels of donor and acceptor where included in the analysis (see Materials and Methods). The ratio of the integral of CFP emission divided by the integral of YFP emission isolated after spectral unmixing of confocal images is proportional to FRET efficiency. Compared to WT channels, the FRET index was significantly reduced for W344R subunits (0.49 ± 0.090 for WT vs 0.10 ± 0.012 for W344R, Figure 3A).

**Figure 3.**
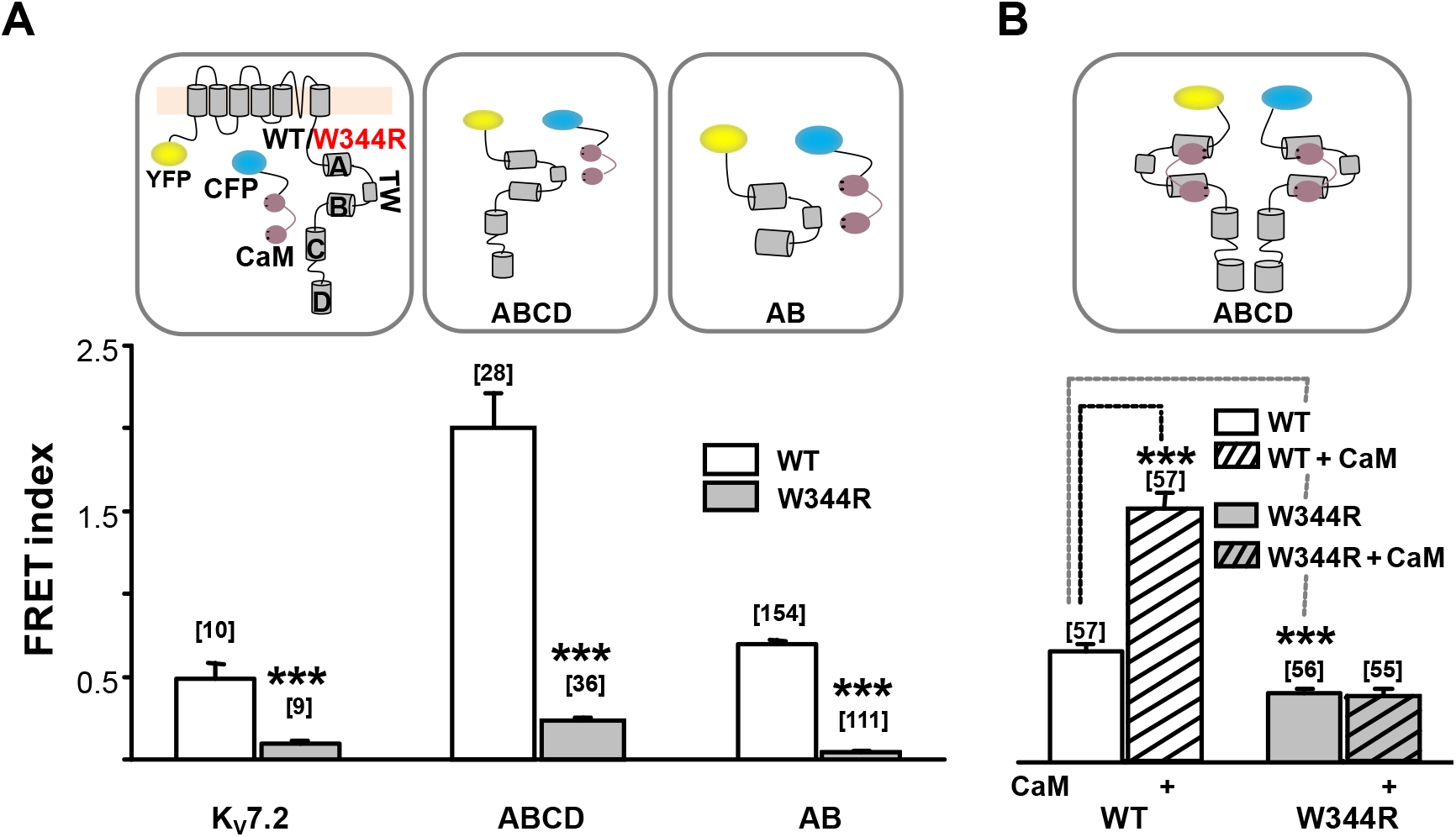
The W344R mutation disrupts K_v_7.2 interaction with CaM in living cells. **A.** FRET index from spectrally unmixed confocal images of HEK293T cells that express YFP-tagged channels or C-terminal domains (AB, ABCD) and CFP-tagged CaM for the indicated configurations. **B.** FRET index between ABCD domains before and after (dashed columns) CaM co-expression. Number of cells is indicated in brackets. ***p < 0.001.

Four alpha helices designated A through D can be recognized within the intracellular C-terminal domain of every K_v_7 channel [16; 37]. CaM embraces the hairpin formed by helices A and B [14; 38; 39]. The CRD is followed by two alpha helices, C and D, which run perpendicular to the membrane [17; 40]. Assembling as either heterotetramer or homotetramer depends on the identity of helix D [37].

To test if the W344R mutation decreases CaM binding to helices A and B of K_v_7.2 *in cellulo*, CFP-CaM was co-expressed with YFP-tagged C-terminal tail containing helices A through D that forms tetrameric complexes (YFP-ABCD), or with YFP-tagged helices A and B (YFP-AB) that forms monomeric complexes [34]. The transfer of energy was significantly larger when the acceptor was ABCD than when it was AB, and, importantly, the FRET index was almost abolished for the W344R mutant proteins (Figure 3A).

We have previously observed FRET in cells expressing helices ABCD (CFP-ABCD + YFP-ABCD) which is related to the ability of the D segment to form tetrameric coiled-coil assemblies [34]. The ABCD resembles a flower bouquet with the coiled-coil helix D corresponding to the pedestal [14]. The transfer of energy between these proteins increases with CaM overexpression (Figure 3B), suggesting that CaM promotes a rearrangement of helix A within the ABCD region [34]. In contrast, the presence of the W344R mutation completely obliterated the FRET response to increased CaM expression, and this FRET response was lower under basal or elevated CaM conditions (Figure 3B), suggesting that the C-terminal region adopted a more relaxed configuration due to the W344R mutation. These results reinforce the proposal that this mutation prevents CaM binding to the CRD of K_v_7.2 channels in living cells.

### The W344R mutation prevents proper folding during translation

We have previously shown that the W344R mutation does not disturb CaM binding to the GST-CRD fusion protein using *in vitro* binding assays, including dansylated-CaM fluorescence emission, Far-Western or Surface Plasmon Resonance [1]. The discrepancy between *in vitro* and *in cellulo* CaM binding could be explained if the polypeptide follows different folding pathways. We hypothesized that the production and isolation of the GST-CRD fusion protein allowed folding *in vitro* of the CRD to a native configuration that could be recognized by CaM. To evaluate this, we used a “folding sensor” (Figure 4A) which has been previously described [15]. This sensor contains helices A and B flanked by mTFP1 and mcpVenus florescent reporters at the protein N- and C-terminal ends, respectively (Figure 4A). Since helices A and B adopt an antiparallel fork configuration, co-expression of CaM brings both fluorophores close to each other, resulting in a FRET index value of 1.70, defined as the ratio of peak emission at 528 nm and 492 nm. Addition of the strong chaotropic agent urea at 6 M reduced of the FRET index to 0.55, which is the same value obtained with mTFP1 alone. Since both mTFP1 and mcpVenus fluorogenic properties are not affected by this treatment, this FRET reduction indicates unfolding of the AB fork (supplemental Figure 4).

**Figure 4.**
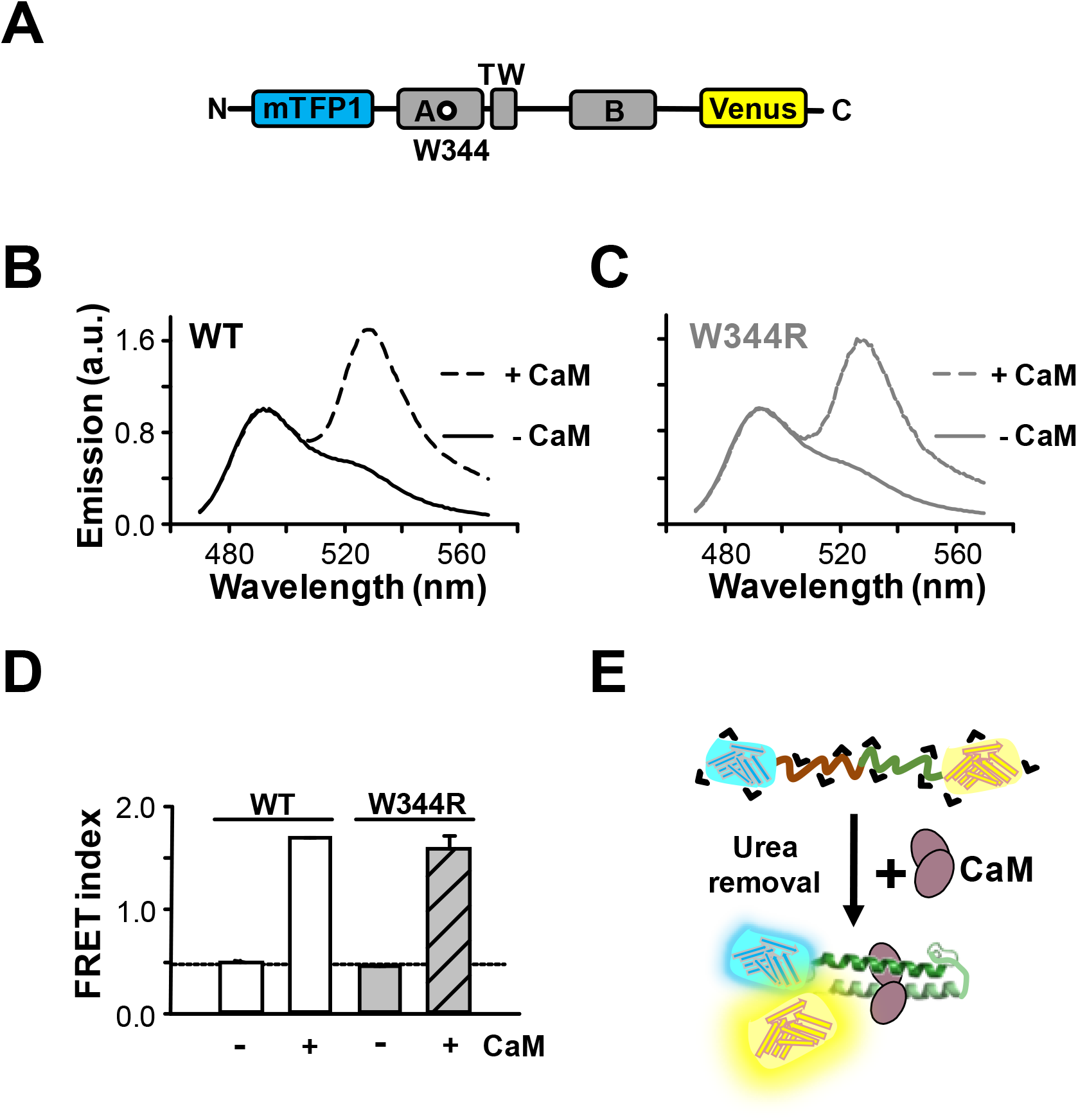
The W344R mutation is compatible with proper folding of the CRD. **A.** Schematic representation of the “folding biosensor”. Helices A and B are flanked by mTFP1 and mcpVenus fluorescent proteins. Emission spectra of WT (**B**) and W344R (**C**) biosensors after denaturalization with 6 M urea and subsequent dialysis without (solid lines) and with (dashed lines) CaM. **D**. FRET index values of WT (white) and W344R (gray) proteins from data as in B and C. The dotted horizontal line marks the index computed in the absence of acceptor. **E**. Schematic interpretation of the experiment: Upon refolding in the presence of CaM, the fluorescent proteins are within FRET distance.

Folding in a cellular context in the absence of CaM was assessed in bacteria, since essential folding mechanisms are shared by prokaryotes and eukaryotes [30], and prokaryotes do not express CaM [41]. Most of the expressed folding sensor molecules were insoluble (Figure 5A, left). The material from inclusion bodies was solubilized with 6 M urea, and thereafter the chaotropic agent was removed by dialysis in the presence or absence of CaM (Figure 4). The emission spectra from reconstituted WT and W344R folding sensors were indistinguishable, presenting FRET index values congruent with proper folding when CaM was present (Figures 4B-E). Thus, the W344R mutation is compatible with the adoption of the native AB fork configuration, consistent with the ability of CaM to bind *in vitro* to the AB protein [1].

**Figure 5.**
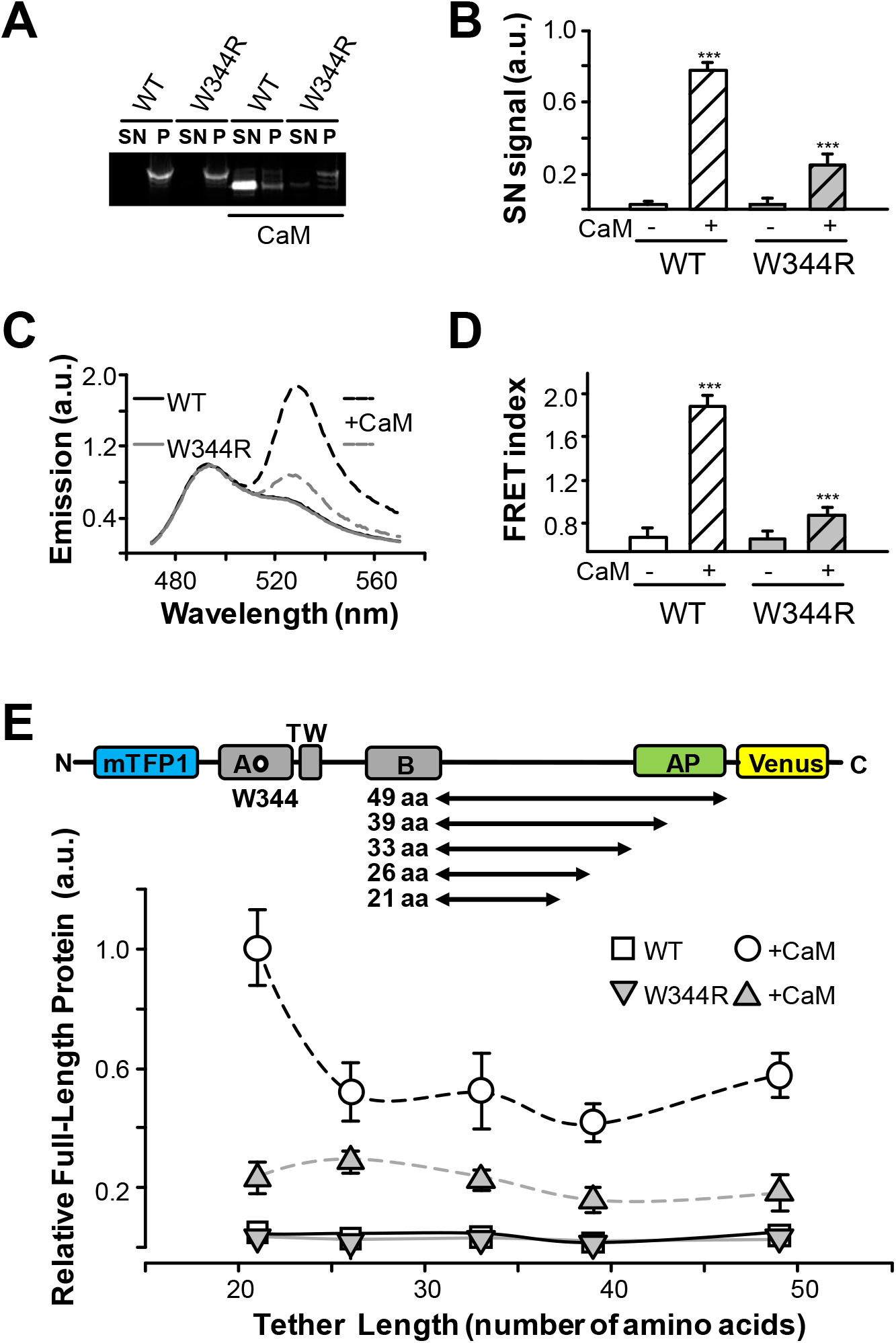
The W344R mutation disrupts folding of the CRD. **A.** Fluorescent image of a pseudo-native SDS-PAGE of bacterial extracts expressing WT or W344R biosensors at 37°C. Proteins were co-expressed (right columns) or not (left columns) with CaM. Soluble (supernatant; SN) and insoluble (pellet; P) protein fractions were separated, and loaded as indicated. **B**. Fluorescence intensity of the supernatant band of SDS-PAGE gels (n = 3). **C**. Emission spectra of the soluble fraction of WT (black lines) and W344R (grey lines) proteins expressed alone (solid lines) or co-expressed with CaM (dashed lines). **D**. FRET index values from spectra as in C. **E.** Top. Schematic representation of the constructs used for *in vivo* cotranslation folding monitoring. The CRD was cloned upstream of the SecM arresting peptide (AP) sequence with tethers of increasing length, ranging from 21 to 49 amino acids from the C-terminal conserved Pro of the SecM AP where translational stalling takes place. mTFP1 and mcpVenus were fused to the N- and C-terminus, respectively. Folding events of the protein domains inside or outside the ribosomal tunnel alleviate SecM stalling and leads to an increase in the ratio of peak emission mcpVenus/mTPF1. Bottom, cotranslational folding profiles of the indicated AB CRD constructs, expressed with and without CaM (n ≥ 4).

The emission spectra of the small soluble fraction of either WT or W344R sensors translated in CaM-free bacteria did not display FRET (Figure 5C), suggesting their unfolded nature. To test if the presence of CaM can induce their proper folding, excess purified CaM was added to the soluble fraction. However, no indication of CaM induced folding was observed as the FRET index remained unaltered after up to 24 hours (supplemental Figure 5), suggesting that the unfolded AB fork was stable and cannot be rescued by *in vitro* addition of CaM.

When CaM was co-expressed in bacteria and therefore it was present during translation, a larger fraction of the WT sensors became soluble (Figures 5A and B) and presented a robust FRET (Figure 5C). In contrast, for the W344R mutant, a large fraction ended up in inclusion bodies (Figures 5A and 5B), although a small soluble fraction displayed significant FRET index (Figure 5D). This FRET index represented 25% of that observed for the WT biosensor, suggesting that there was a small proportion of properly folded sensors carrying the W344R mutation.

To assess how the rate of translation affects the outcome, the same experiments were performed lowering the temperature to 18°C during the induction of translation, but no significant difference was found compared to 37°C (supplemental Figure 6).

We exploited the ability of the SecM translational arresting peptide (AP) to act as a force sensor that detects folding of proteins in the ribosomal exit tunnel during translation [42–45]. The SecM AP interacts with the ribosome tunnel and detains protein synthesis, unless external force acting on the polypeptide chain “pulls-out” the AP, thereby relieving translational arrest [42]. Such pulling force can be induced by cotranslational protein folding [42; 43], with equivalent results *in vitro* and *in cellulo* [44], and identifies the same cotranslational folding transitions as do other methods, such as real-time FRET, photoinduced electron transfer, and NMR [46]. The K_v_7.2 CRD was cloned upstream of the SecM arresting peptide sequence with tethers of increasing length and flanked by mTFP1 and mcpVenus fluorescent reporters in the N- and C-terminus respectively (Figure 5E). In this paradigm, if the protein is stalled, the result is an mTFP1-tagged truncated protein. In contrast, if CaM participates in folding of the CRD during translation and the full-length protein is translated; fluorescent signals from both mTFP1 and mcpVenus could be recorded (see supplemental Figure 7). By measuring the stalling efficiency (as fraction of full-length reporter protein) for a series of constructs of increasing tether length, it is possible to identify when the protein starts to fold during translation [45; 46].

Therefore, we used the ratio of emission at the peak wavelength for mcpVenus and for mTFP1 to assess the fraction of full-length reporter protein (Figure 5E). The ratio was very low when WT or W344R AP reporters were expressed alone, but there was a robust signal for WT when expressed in the presence of CaM. This suggests that CaM exerts a fundamental role for CRD folding during translation. The index decreased as the length of the tether increased, reminiscent of profiles described for other proteins [44]. Although lower than WT, CaM also increased the translation of full-length W344R reporter, suggesting that a fraction of the W344R mutant reporter was folding during translation when CaM was present. These results are concordant with FRET observed for the AB sensor (Figures 5C and 5D).

### Two orientations for tryptophan and arginine at position 344

To explore the stability of the W344R CaM/K_v_7.2-AB complex *in silico*, binding affinities were derived using the Rosetta Flex ddG software. Random changes in the backbone angles near the site of interest are introduced, generating a trajectory of structures which are classified with respect to their resulting energy values. Afterwards, each of these random structures are ruled out or accepted, according to a Metropoli Montecarlo criteria, and resulting in small winning population of the energetically most stable configurations.

We found that in most of the cases the side chain of tryptophan at position 344 is oriented towards CaM in the structures of the AB/CaM complex of different K_v_7 subunits [14; 15; 20; 38; 40]. Interestingly, detailed visualization of the final accepted trajectory snapshots of Rosetta moves revealed a small population (1.6% for WT and 3.4% for W344R) in which the side chain became tilted (T) towards the rest of the A helix, instead of targeting the CaM C-lobe, as it appears in the native (N) WT structure (Figure 6A).

**Figure 6.**
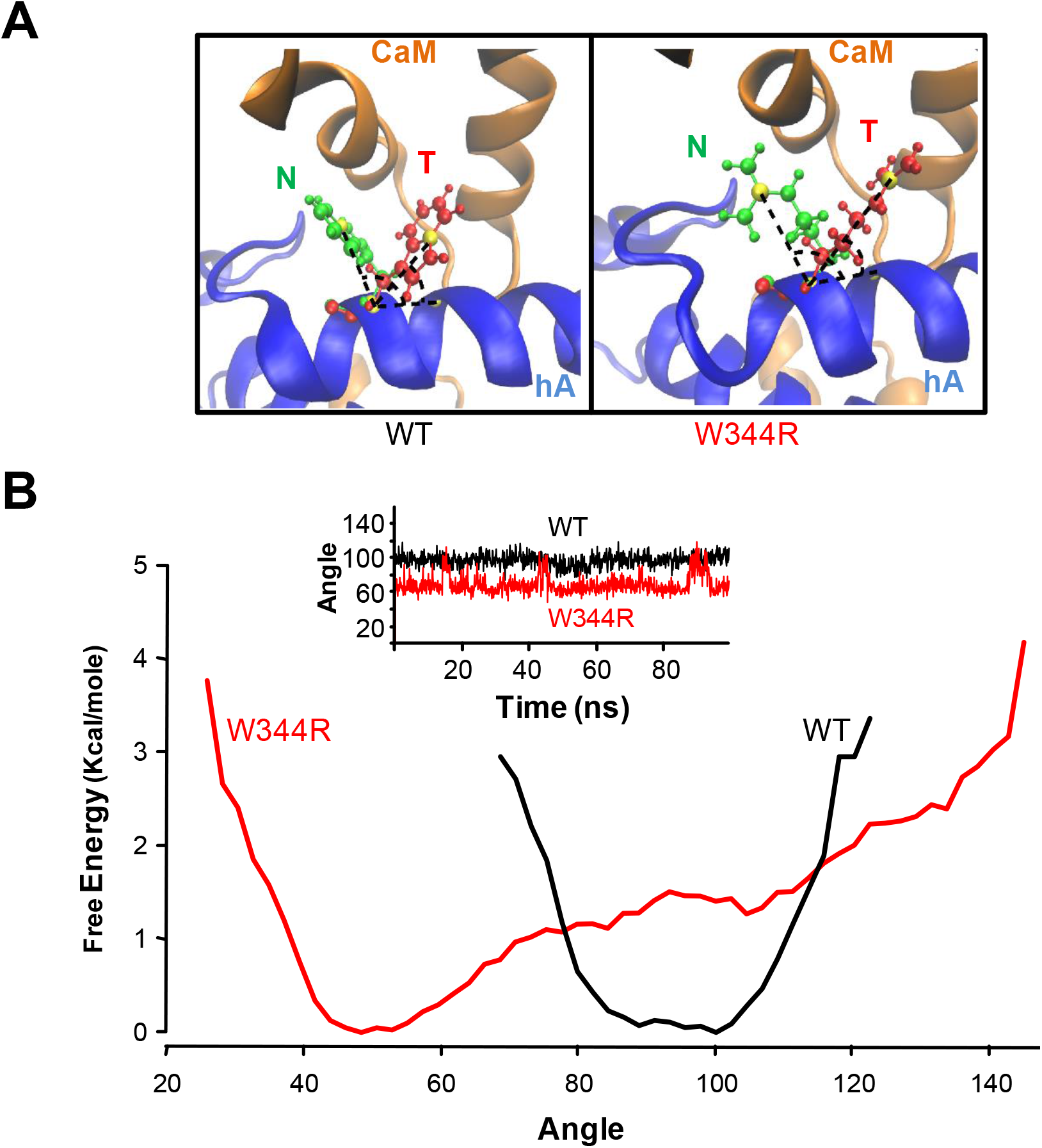
Two orientations for the lateral chain at position 344. **A.** Overlaid representation of both Tilted (T, red) and Native-like (N, green) structures of WT (left) and W344R mutant (right), where hA is blue whereas the CaM C-lobe is orange. Angles between hA and T or N structures are indicated. **B**. Projection of the free energy in the angles between hA and tryptophan or arginine at position 344. Note that the angles of the N and T orientations depend on the presence or absence of CaM. Due to the mathematical computations involved, as WT is not able to explore angles in the ranges (20°-65°) and (125°-150°), the resulting free energy would be infinite, and it is not showed. The inset shows the time series of the angle through the simulation (Supplemental Figure 9).

The computed value of the *ΔΔG* for W344R was 5.1 kcal/mol. This is a common value of a destabilizing mutation [5; 47; 48](supplemental Figure 8), suggesting that CaM binding to W344R should be disrupted, which seems inconsistent with the actual *in vitro* binding data (supplemental Figure 8). To unravel this inconsistency, we next computed all-atom energy.

To elaborate a hypothesis on the mechanism underlying the differential effect of the W344R mutation *in vitro* and *in cellulo*, we determined which orientation is the most stable in absence of CaM. For this purpose, all-atom molecular dynamics (MD) simulations of the hA-TW-hB segment not engaged with CaM in a water cubic box for both WT and mutant were performed. After equilibration, the tryptophan residue at position 344 converges to the N position, whereas the arginine residue adopted the T orientation, and subsequent MD simulations were computed from these starting positions, respectively (Figure 6B). In WT, tryptophan remains stable in the N configuration during 100 ns, and no transitions to the T state were observed. Furthermore, when W344 was forced to start from the T configuration, it transited to the N state in a few ns and remained there, confirming that the N configuration is more stable (see supplemental Figure 9). The energy needed to overcome the conformational change of the side chain from N to T is higher for the WT, which may underlie its inability to explore the T configuration. In W344R mutant, transitions between the two states were observed for the arginine residue, remaining more time in T (Figure 6B), suggesting that the energy barrier between the two states in the absence of CaM is smaller. Since W344R remains more time in T than in N, we conclude that the T configuration is the most stable when not engaged to CaM. To represent this information from an energetic perspective, the potential of mean force for the angles formed by the side chain and backbone were computed using the weighted histogram analysis method [49]. Figure 6B shows that the WT has a defined potential well around the N configuration. Conversely, W344R has its lowest energy configuration in T and the potential barrier between the two states is lower, so that it is more common to find the T configuration but the N is not energetically banned.

To identify which configuration of the mutated K_v_7.2 channel favors binding to CaM, the average of the non-bonding interaction energy between the residue W344R and CaM was computed using the CHARMM36 all-atom energy function for all the snapshots of Rosetta trajectories. In W344R mutant, the energy for T configuration was −16 kcal/mol, and that for the N configuration was −76 kcal/mol. In WT, the computed energies were −10 kcal/mol and −20 kcal/mol for the T and N configurations, respectively. These results suggest that the N configuration is more compatible with CaM binding than the T configuration, and that arginine is more stable than tryptophan when helix A is in complex with CaM. This may explain the ability of both WT and W344R to bind CaM *in vitro* (see supplemental Figure 8).

## DISCUSSION

The analysis of mutations at helix A of the CRD of K_v_7.2 subunits has revealed an obvious relationship between CaM binding and function: subunits that do not bind CaM *in vitro* and *in cellulo* are retained at the ER, and channel function is concomitantly abolished [2; 13; 32; 33](supplemental Figure 1).

The W344R mutation found in patients with hereditary epilepsy [1; 31] is unique because it does not affect CaM binding to the CRD of K_v_7.2 *in vitro* [1](supplemental Figure 8), but abolish current and surface expression of K_v_7.2 channels (Figures 1–2). Two main hypotheses could explain this striking deviation from the common trend. One possibility is that this mutation may lock the gate of the channel in a closed state, for example, by impeding the expansion of the inner gate formed by the S6 bundle crossing that is directly attached to helix A. Alternatively, this mutation may prevent CaM binding within the cellular environment, but not in *in vitro* assays (supplemental Figure 8).

Here, we show that the W344R mutation prevents FRET between CaM and K_v_7.2 subunits in cells. In addition, the tetrameric assembly of the four A-D helices in mutant K_v_7.2 [34] becomes insensitive to CaM abundance (Figure 3). Therefore, the W344R mutation disrupts CaM binding to the isolated CDR or the full-length channel in living cells. Importantly, our FRET sensor that detects folding of the antiparallel AB fork, and the assessment of release from stalled ribosomes, revealed that the W344R mutation disrupts CRD folding during translation in the presence of CaM *in cellulo* (Figures 4–5), indicating that the majority of the W344R mutant proteins fail to adopt a properly folded state during translation in living cells. Our results indicate that the presence of CaM during translation can be considered essential for proper folding of K_v_7 CRD, in agreement with previous proposals [50; 51].

What are the differential properties between tryptophan and arginine that lead to the failure of W344R to adopt a configuration that can be recognized by CaM *in cellulo*? We discovered two possible orientations (N and T) for the residue at position 344 (Figure 6). When the CRD is not in complex with CaM, the adoption of the T orientation in WT CRD is very rare, concordant with atomic models [14; 15; 20; 38; 40], whereas the T configuration is stable and preferred over the N configuration for W344R CRD (Figure 6). However, when helix A is in complex with CaM, the local interactions with the arginine residue in the W344R mutant are more favorable than the tryptophan residue in the WT (Figure 6), consistent with *in vitro* binding data that demonstrates similar CaM interaction with WT and W344R CRD [1](see supplemental Figure 8).

We speculate that the vectorial nature of translation of the mutated K_v_7.2 channel, with the subsequent decrease on its available configurational space, could favor the T configuration of the W344R side chain during the transit of the nascent chain in the ribosomal tunnel. Consequently, after the nascent chain emerges from the ribosome in a eukaryotic cellular environment with abundance of CaM molecules, the W344R residue might remain in the metastable T configuration, which could prevent the proper binding between CaM and the mutated channel. In addition, this configuration may alter interactions with molecular chaperones or favor degradation. On the other hand, *in vitro* experiments might enlarge the accessible configurations for the W344R side chain, allowing it to find the most stable N configuration when interacting with CaM, favoring the mutated K_v_7.2 channel to properly bind CaM. The key difference is that *in vivo* the N-terminal portion can start folding before the C-terminal portion has been synthesized or is still within the ribosomal tunnel. In contrast, refolding of the full CRD *in vitro* can begin via interactions anywhere along the peptide chain [52]. In summary, the adoption of the T orientation favored by the W344R mutation may represent a kinetic trap in the path to a native fold [53], although more data is required to fully understand the process.

Genetic protein folding defects could be segregated into two categories: those due to mutations that are unsuited to adopt the native fold and those that can adopt the native structure with the use of chemical chaperons, low temperature or ligands. For instance, the function of the N258S K_v_7.2 mutation, located in the extracellular turret just after S5, can be partially recovered with K_v_7.3 subunits, by culturing cells at lower temperatures or in the presence of the K_v_7-binding drug retigabine [28]. This second group could be divided further into those defects that lead to a weaker or unstable structure, and those that are compatible with a stable native configuration but fail to find the proper folding pathway. The W344R mutant fits readily in this latter category. We are not aware of any other pathological mutation with these properties and, therefore, this mutant may be the first representative within this group.

A remaining question is when and where does the W344R mutant deviate from the proper folding pathway. Some nascent chains can adopt an alpha helix configuration in the restricted space of the ribosomal tunnel, and can fold into tertiary structures in its vestibule [30]. Analysis of the amino acid sequence of helix A using the Agadir server predicts that both WT and W344R will fail to form a stable alpha helix in solution, with scores of 0.4 and 0.7 (in a 0 to 100 scale), respectively. However, even unstable peptides may form helical structures in some regions of the ribosomal tunnel [54–56]. The vectorial nature of protein synthesis, the spatial constrains and physicochemical properties of the ribosomal tunnel, can guide the folding trajectory of the nascent peptide [30]. Our data and other observations are consistent with the idea that the emerging helix A segment, located at the N-terminus of the CRD, starts to fold before the C-terminal part of the protein is synthesized [57], and will initiate interactions with the CaM C-lobe in the vestibule or outside the ribosome during translation. It is not known if the emerging segment is already folded, or if CaM induces the adoption of an alpha helix. Our data is compatible with the idea that CaM fails to promote alpha helix formation to the mutant W344R nascent chain, whereas *in vitro* binding is best described by selection of the properly folded molecules [58–60].

In summary, we show here that the autonomously folding calcium responsive domain (CRD) carrying the W344R mutation is not recognized by CaM in a cellular *in vivo* context, but it does so after refolding *in vitro*. This mutation impedes proper folding during translation within the cell by forcing the nascent chain to follow a folding route that leads to non-functional ion channels. Thus, although it carries all the information for the native 3D configuration, it fails to reach it *in vivo*. Our MD simulations suggest a reasonable hypothesis for the underlying mechanism. Thus, this study provides a mechanistic insight into co-translational folding defects, which may represent a widespread mechanism that contributes to pathophysiology.

## Author Contributions

A.V. conceived the study and participated in its design and coordination. J.U., A.A., C.G-P., and A.M-M. carried out experiments, contributed to figure preparation, and manuscript preparation. O.R.B., A.M-M., A.B. and A.L. performed *in silico* analysis, contributed to figure preparation, and manuscript preparation. J.Z. and H.J.C. contributed to neuronal characterization, figure preparation, and manuscript preparation. E.N. and C.M. prepared reagents, and contributed to experimental design and figure preparation. All authors read and approved the final manuscript.

## Data and material availability

Data and material are available upon request.

## SUPPLEMENTAL MATERIAL

## MATERIAL AND METHODS

### CELL CULTURE AND TRANSFECTION

Human Embryonic Kidney cells (HEK293T) were cultured in DMEM (Dubelcco’s modified Eagle’s medium), supplemented with 10% of fetal bovine serum, 1% no-essential amino acids and 1% of gentamicin. Cell cultures were maintained in 5% CO_2_ at 37°C. Cells were transiently transfected with desired cDNAs using Polyethylenimine (1 μg/μl; Polysciences) for electrophysiological studies, FRET and Co-immunoprecipitation experiments.

### ELECTROPHYSIOLOGY

The human isoform 3 K_v_7.2 (Y15065) and K_v_7.3 (NM004519) cDNA. The subunits were tagged at the N-terminus with mCFP or mYFP fluorescent proteins and cloned into pcDNA3.1, respectively were used in this study. N-terminal tags have no impact on the electrophysiological properties of the expressed channels [7; 27]. Macroscopic currents were recorded at room temperature (22°C) in the whole-cell configurations of the patch-clamp technique using HEKA patch clamp EPC8. Borosilicate capillary glass (Sutter instrument) was pulled obtaining a tip resistance of 1-3 MΩ after filled with the internal solution. This solution contains (in mM): 125 KCl, 10 Hepes (K), 5 MgCl_2_, 5 EGTA, 5 Na_2_ATP, adjusted to pH 7.2 with KOH and the osmolarity adjusted to ~300 mOsm with mannitol.

Following patch rupture, whole-cell membrane capacitances were measured from integration of the capacitive transients elicited by voltage steps from −50 to −60 mV, which did not activate any time dependent membrane current. Series resistances were compensated 80% in order to minimize voltage errors and were checked regularly throughout the experiment to ensure that there were no variations with time. The voltage-clamp experimental protocols were controlled with the “Clampex” program of the “pClamp” software (Molecular Devices). HEK293T cells were perfused with the external solution containing (in mM): 140 NaCl, 4 KCl, 2 MgCl_2_ (6 H_2_O), 10 Hepes-Na, 2 CaCl_2_ and 5 glucose, pH 7.4 with NaOH and the osmolarity adjusted to ~320 mOsm with mannitol.

The amplitude of the K_v_7 current was defined as the peak difference in current relaxation measured at −30 mV after 500-1,500 ms pulses to −110 mV (all channels closed) and to +20 mV (all channels opened).

### CO-IMMUNOPRECIPITATION (Co-Ip)

A confluent T-75 flask of HEK293T cells were co-transfected with 5 μg of KCNQ2-WT cDNA in pcDNA3.1, N-terminally tagged with either CFP or Myc, and 5 μg of CFP-KCNQ2-W344R cDNA for Co-Ip experiments to analyze protein-protein interaction. Twenty four hours after transfection, HEK293T cells were solubilized for 30 min at 4°C in RIPA buffer, containing (mM): 20 Tris-HCl (pH 7.5),150 NaCl, 5 EDTA, 1% NP40 and protease inhibitors (1X Complete; Roche Applied Science). The lysate was centrifuged at 800 g for 15 min and the insoluble material was removed after centrifugation at 13,000 g for 15 min, after which the lysate was pre-cleared for 1 h at 4 °C with 40 μl of equilibrated protein G-Sepharose beads (GE Healthcare). The day before, anti-*c*-Myc (Sigma-Aldrich) antibody was immobilized overnight at 4°C with 40 μl of equilibrated protein G-Sepharose beads and washed six times with RIPA buffer. Precleared lysates were incubated overnight at 4°C with protein G-antibody mix. After 6-8 washes with RIPA buffer, the immunoprecipitated proteins were released by heating at 90°C 5 min in SDS sample buffer and were probed with anti K_v_7.2 antibody.

### WESTERN BLOT

Immunoprecipitation samples were fractionated on 6% or 15% SDS-polyacrylamide gels and transferred to polyvinylidene fluoride membranes (PVDF) (Millipore). Membranes were blocked in TBS solution (0.05% Tween-20 in PBS containing 5% milk). Then, they were incubated with the monoclonal primary antibody: anti-K_v_7.2 (1:1,000; Neuromab). Secondary antibody was goat anti-mouse IgG horseradish peroxidase conjugate (1:5,000; Bio-Rad). Blots were developed using the Luminata Forte Western HRP substrate reagent (Millipore) and images were digitalized with a Thermo Scientific MYECL Imager. Densitometry of the bands was measured by FIJI software. It was calculated dividing co-immunoprecipitated protein by immunoprecipitated protein, i. e. CFP-K_v_7.2/c-Myc-K_v_7.2.

### FRET IN LIVING CELLS

Cells were plated at ~60% confluence onto 30-mm round coverslips in six-well plates. Cells were transfected as described above. For CFP-CaM binding to YFP-K_v_7.2, YFP-ABCD or YFP-AB the transfection ratio used was 1:5 with a total of 0.6 μg DNA per M35 dish. For assembly experiments, the ratio was 1:1:2 (0.5 μg of each FCP-tagged ABCD and 1 μg of CaM or 1 μg of empty pcDNA3.1 his/c-Myc vector). Twenty-four hours after transfection, coverslips were placed in an imaging chamber and imaged maintaining them in buffer solution composed of (mM): 140 NaCl, 5 KCl, 1 MgCl_2_, 2 CaCl_2_, 10 glucose and 10 Na-Hepes, pH 7.4 at room temperature.

Images were recorded using a Nikon D Eclipse TE2000-U fluorescence microscope (Nikon Instruments, Tokyo, Japan) equipped with a confocal scanning head and a spectral detector module. Images were captured using a 60X oil objective, with the pinhole opened (150 μm) and using the 405 nm laser line (Coherent, Santa Clara, CA, USA) or the 488 nm line (Melles-Griot, Rochester, NY, USA) for direct CFP or YFP excitation, respectively. To assure homogeneity in donor an acceptor expression, cells displaying a signal with emission values between 20 and 170 (arbitrary units) when excited at 405 nm (to record FRET signal) and when excited at 488 nm (to record acceptor emission only) were included in the analysis with FIJI.

The spectral detector allows simultaneous recording of 32 images, each registering a 5 nm band of the spectrum, covering 450–610 nm. After spectral unmixing with EZ-C1 Nikon software, using cells expressing CFP or YFP alone as reference, as described previously [34], the area under the spectra was measured and a FRET index was calculated as FRET index = YFP_405_/CFP_405_, where YFP is the integral of emission signal for YPP_405_, and CFP_405_ is the integral of the emission signal for CFP after excitation with the 405 nm laser line.

### EXPERIMENTAL ANIMALS AND NEURONAL CULTURE

All procedures involving animals were reviewed and approved by the Institutional Animal Care and Use Committee at the University of Illinois Urbana-Champaign in accordance with the guidelines of the U.S National Institutes of Health (Protocols 15222). Primary dissociated hippocampal cultures were prepared from 18-day old embryonic rats and transfected with plasmids (total 0.8 μg) at 5 DIV as described [32].

### IMMUNOCYTOCHEMISTRY

Primary dissociated hippocampal cultures were prepared from 18-day old embryonic rats, transfected with plasmids (total 0.8 μg) at 5 DIV, and immunostaining for surface and total K_v_7 subunits and Axonal Initial Segment (AIS) markers in hippocampal neurons were performed at 48 h post transfection as described [32]. Fluorescence images were acquired as described [5; 32] using a Zeiss Axio Observed inverted microscope equipped with a Zeiss AxioCam 702 mono Camera and ZEN Blue 2.6 software, and stored with no further modification as CZI and 16-bit TIFF files. Within one experiment, the images were acquired using the same exposure time to compare the fluorescence intensity of the neurons transfected with different constructs.

The background subtracted mean fluorescence intensity of the soma, the axon within 0-30 μm of the beginning of the axon (AIS), the axon between 50 and 80 μm from the beginning of the axon (distal axon), and the major primary dendrites were quantified using ImageJ Software (National Institutes of Health) as described [32].

### TRANSLATION ANALYSIS

The DNAs cloned in pProHex-HTc corresponding WT and W344R CRD flanked by mTFP1 and mcpVenus fluorophores in the N- and C-termini respectively were transformed in *E. coli* BL21 cells alone or together with the pOKD4 plasmid carrying the CaM gene. Cells were grown over night at 37°C and diluted into 20 ml of fresh LB for further growing at 37°C till OD_600_ 0.6. Protein expression was induced during 3 h at 37°C or overnight at 18°C by addition of 1 mM IPTG. Cells were harvested by centrifugation at 7,000 rpm for 10 min. The cell pellets were resuspended in lysis buffer 50 mM Hepes, pH 7.4, 120 mM KCl, 5 mM NaCl, 5 mM EGTA, 1 mM DTT and protease inhibitors (1X Complete; Roche Applied Science), and similar OD values were fitted for all the samples. The cellular cultures were sonicated 3 times, 5 s ON, 5 s OFF and centrifuged at 19,000 x g during 30 min for supernatant and pellet separation. The pellets were resuspended in the same buffer volume used before. Protein solubility was studied by SDS-PAGE electrophoresis (10%), by analyzing the protein amount present in the same volume of pellet and supernatant fractions. The gels were visualized using Versadoc imaging equipment, exciting using blue or green leds, combined with 530BP28 or 605BP35 emission filters. The protein amount in the pellet and in the supernatant was estimated relative to the total protein amount by quantification of the gel bands using the ImageJ software. The protein soluble fractions were also analyzed in a Fluoromax-3 fluorimeter by recording the emission spectra of mTFP1 and mcpVenus fluorescent proteins upon excitation at 458 and 515 nm respectively.

### UREA MEDIATED PROTEIN DENATURALIZATION AND RENATURALIZATION

K_v_7.2 WT and W344R cloned in pProHex-HTc were transformed in *E. coli* BL21 cells in the absence of CaM and grown over night at 37°C. Cell cultures were diluted into 20 ml of fresh LB for further growing at 37°C till OD_600_ 0.6. Protein expression was induced during 3 h at 37°C by addition of 1 mM IPTG. Cells were harvested by centrifugation at 7,000 rpm for 10 min and resuspended in 500 μl lysis buffer 120 mM KCl, 50 mM Hepes, pH 7.4, 1 mM PMSF, 1% Triton and protease inhibitors. The cellular cultures were sonicated 3 times, 5 s ON, 5 s OFF and centrifuged at 19,000 x g during 30 minutes for supernatant and pellet separation. The pellets were resuspended in 120 mM KCl, 50 mM Hepes, pH 7.4, and protease inhibitors (buffer A) and centrifuged at 19,000 x g for 30 minutes. The pellets were resuspended in buffer A supplemented with 1 M urea and incubated for 20 minutes. Samples were centrifuged again at 19,000 x g for 30 minutes and resuspended in buffer A containing 6 M urea for protein extraction. After 30 minutes incubation at 4°C, the solubilized proteins were collected by centrifugation at 19,000 x g during 30 min and diluted to 0.1 mg/ml. Diluted proteins were dialyzed in the absence and presence of CaM against buffer 120 mM KCl, 50 mM Hepes, pH 7.4, 5 mM DTT and urea decreasing concentrations. Proteins were finally dialyzed in 120 mM KCl, 50 mM Hepes, pH 7.4, 5 mM NaCl, 5 mM EGTA and 5 mM DTT for further fluorescence recordings.

### ARRESTING PEPTIDE EXPERIMENTS

The sequence coding for the SecM arresting peptide (AP) FSTPVWISQHAPIRGSP was inserted between helix B and mcpVenus into the DNAs cloned in pProHex-HTc corresponding to WT and W344R CRD flanked by mTFP1 and mcpVenus fluorophores in the N- and C-termini respectively. The sequence (EFYVGYVPGGSPGRPGGSRPHVGSGGQQGSHV) of the linker joining helix B with SecM AP contained restrictions sites that allowed creating a library with 21, 26, 33, 39 and 49 amino acids. The constructs were transformed in *E. coli* BL21 cells alone or together with the pOKD4 plasmid carrying the CaM gene. Single colonies were used to start overnight cultures, which were induced and processed as described in translation analysis. The protein soluble fractions were also analyzed in a Fluoromax-3 fluorimeter by recording the emission spectra of mTFP1 and mcpVenus fluorescent proteins upon excitation at 458 and 515 nm respectively.

### STABILITY CALCULATIONS

Binding affinities have been computed for five different mutations at position 344 using the *Rosetta Flex ddG* [47] prediction protocol for the CaM/K_v_7.2-hAB complex (PDB: 6FEG [15]). In short, WT and mutant models are generated by performing random displacements of the protein backbone named “backrub moves” in a shell of 8 Å around the mutation site. Side chains of both WT and mutants are optimized by assigning a score to each mutation, based on the all atom Rosetta Energy Function 2015 [48]. The resulting score associated to the mutation is compared with the WT to compute *ΔΔG* = *ΔG_Mutation_ - ΔG_WT_*. Therefore, if *ΔΔG* > 0 the WT would show a stronger binding affinity compared to the mutated one, and the opposite for *ΔΔG* < 0. Following Rosetta’s protocol, fifty different simulations were performed for each mutant, and each one consists of 50,000 backrub moves. These random moves are accepted or rejected based on the Metropolis criterion with an energy of 1.2 kT, and the final value of the binding affinity is the mean value of all 50 simulations. The backrub moves constitute a random trajectory in the angles and provides a sampling of possible configurations of the residue 344 side chain. For each move the free energy difference (*ΔG*) is calculated considering the Rosetta energy function 2015 and its final value is the mean value of all 50 simulations.

Additionally, as atomic coordinates are saved every 5,000 steps and each simulation generates 10 snapshots of the trajectory, we used the VMD interface together with the CHARMM36 all-atom force field [61] to analyze in detail the structural characteristics of the simulations.

Finally, molecular dynamics simulations were carried out of the WT and mutant helices A, TW and B with the software NAMD 2.13 and the CHARMM36 all-atom force field. The input structure was PDB 6FEG [15] eliminating the N-terminal residual amino acids up to residue number 328. The simulation was performed in a periodic cubic box of TIP3P water so that the minimum distance of any protein atom and the edge of the box was at least 6.1 Å. A concentration of 120 mM of KCl and 5 mM of NaCl was introduced to mimic neuron physiology. SHAKE bond length constraints were applied to all bonds, nonbonded interactions were calculated by the particle-mesh Ewald method with a cutoff of 12 Å. The simulation was first minimized using 1,000 steepest descent steps, after that, 0.5 ns were simulated in the canonical ensemble at 298 K, keeping the same temperature using Langevin dynamics with a Langevin damping of 0.5 ps^-1^. After that, 1.5 ns of NPT ensemble was simulated in order to accommodate the periodic cell and avoid the formation of vacuum bubbles in the solvent with a pressure target of 1 atmosphere. Finally, 100 ns were simulated in NPT ensemble for both WT and W344R mutant systems.

### STATISTICAL ANALYSIS

Values are presented as the mean ± SEM. The differences between the means were evaluated using the unpaired Student’s t-test or ANOVA with Mann-Whitney post hoc on SigmaStat Statistic (Sigmaplot 11), where values of *p* < 0.05 were considered significant. The number of cells in each experiment is indicated in brackets in the figures. The results are from two or more independent batches of cells. In all figures *, **, and *** indicate significance at *p* < 0.05, *p* < 0.01, and *p* < 0.001, respectively.

Images from hippocampal neurons were analyzed using Origin 9.1 (Origin Lab), the Student’s t-test and one-way ANOVA with post-ANOVA Tukey’s and Fisher’s multiple comparison tests were performed to identify the statistically significant difference with a priori value *p* < 0.05 between two groups and for > three groups, respectively.

---

**Supplemental Figure 1.**
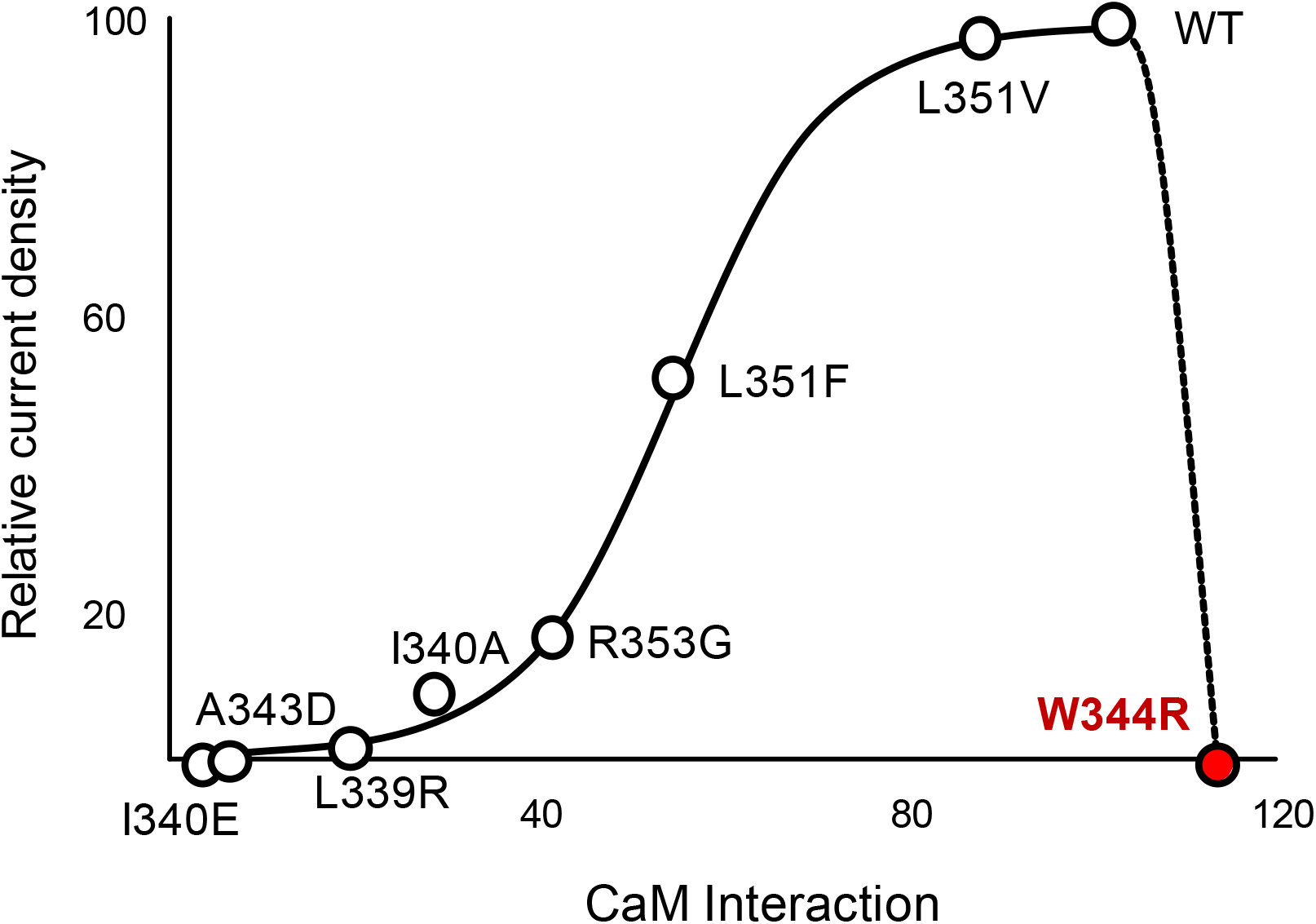
Relationship between current densities of K_v_7.2 channels carrying the indicated mutations in the helix A segment co-expressed with K_v_7.3 subunits and binding to CaM. The W344R mutant deviates from the relationship. Adapted from [1; 2].

**Supplemental Figure 2, related to Figure 2.**
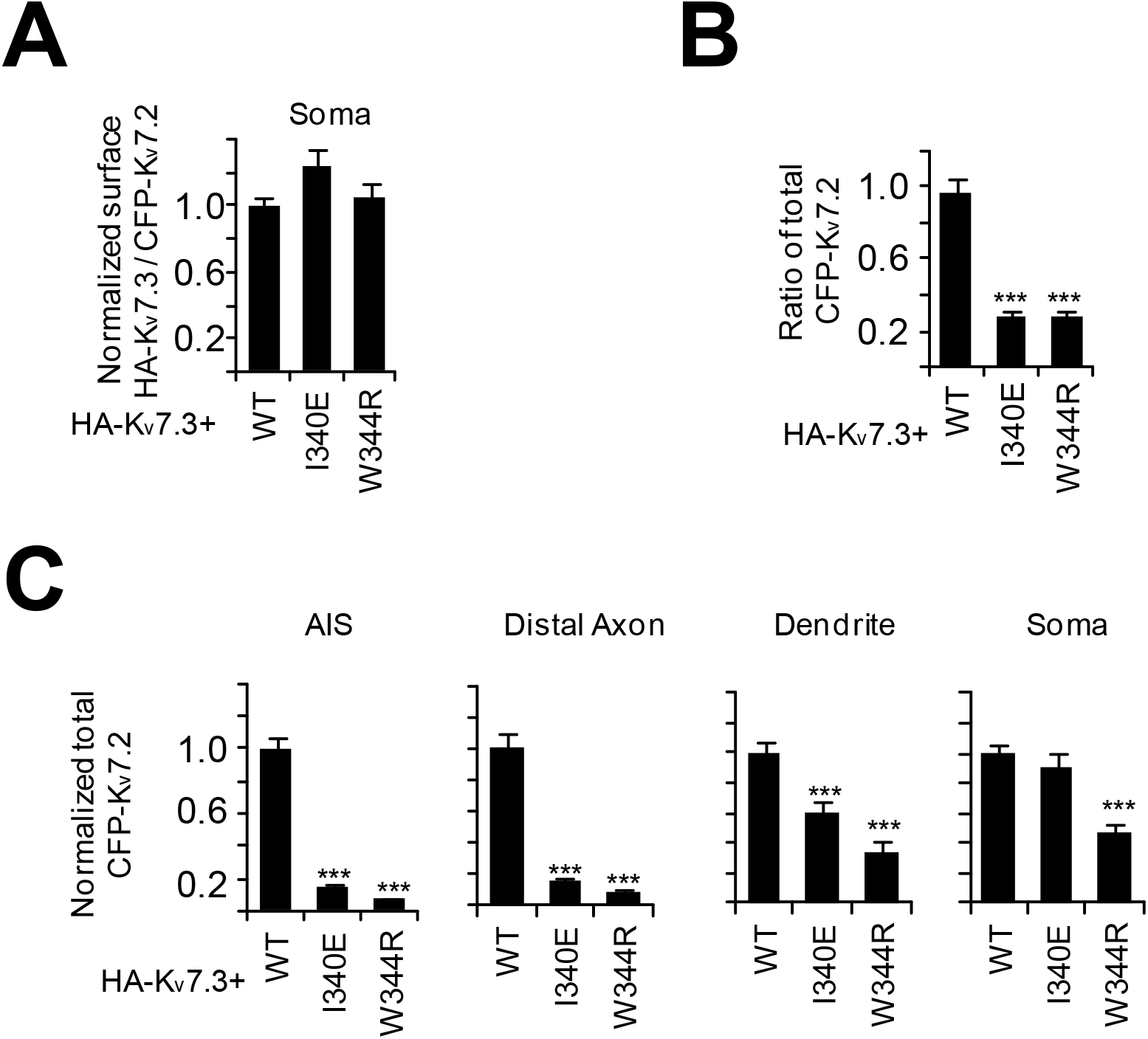
I340E and W344R mutants severely reduced surface and total expression of heteromeric HA-K_v_7.3/CFP-K_v_7.2 in the axons of cultured hippocampal neurons. **A.** The Axon/Dendrite ratio was computed for surface HA-K_v_7.3/CFP-K_v_7.2 in the soma. **B-C** Total CFP-K_v_7.2 fluorescent intensities. Sample numbers are: WT (n = 19), I340E (n = 14), and W344R (n = 13). Data represent Ave ± SEM. One-way ANOVA with post-hoc Fisher’s LSD test was conducted. ***p < 0.005.

**Supplemental Figure 3, related to Figure 2.**
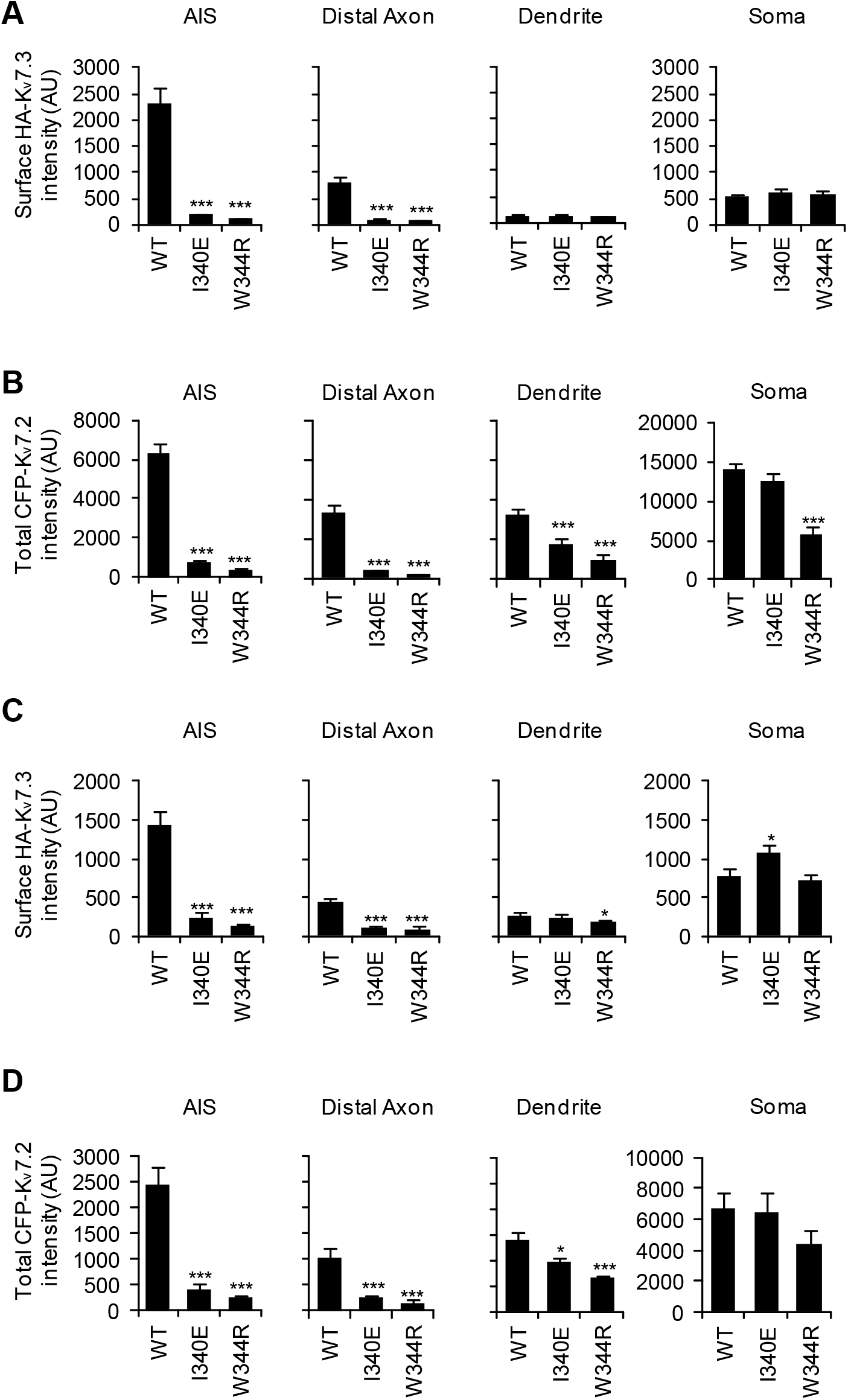
Background-subtracted fluorescent intensities of surface HA-K_v_7.3 (**A, C**) and total CFP-K_v_7.2 (**B, D**) from two individual experiments: experiment 1 (A-B), experiment 2 (C-D). Sample numbers are: (A-B) WT (n = 14), I340E (n = 9), and W344R (n = 10); (C-D) WT (n = 5), I340E (n = 5), and W344R (n = 3). Data represent Ave ± SEM. One-way ANOVA with post-hoc Fisher’s LSD test was conducted. *p < 0.05 ***p < 0.005.

**Supplemental Figure 4, related to Figure 4.**
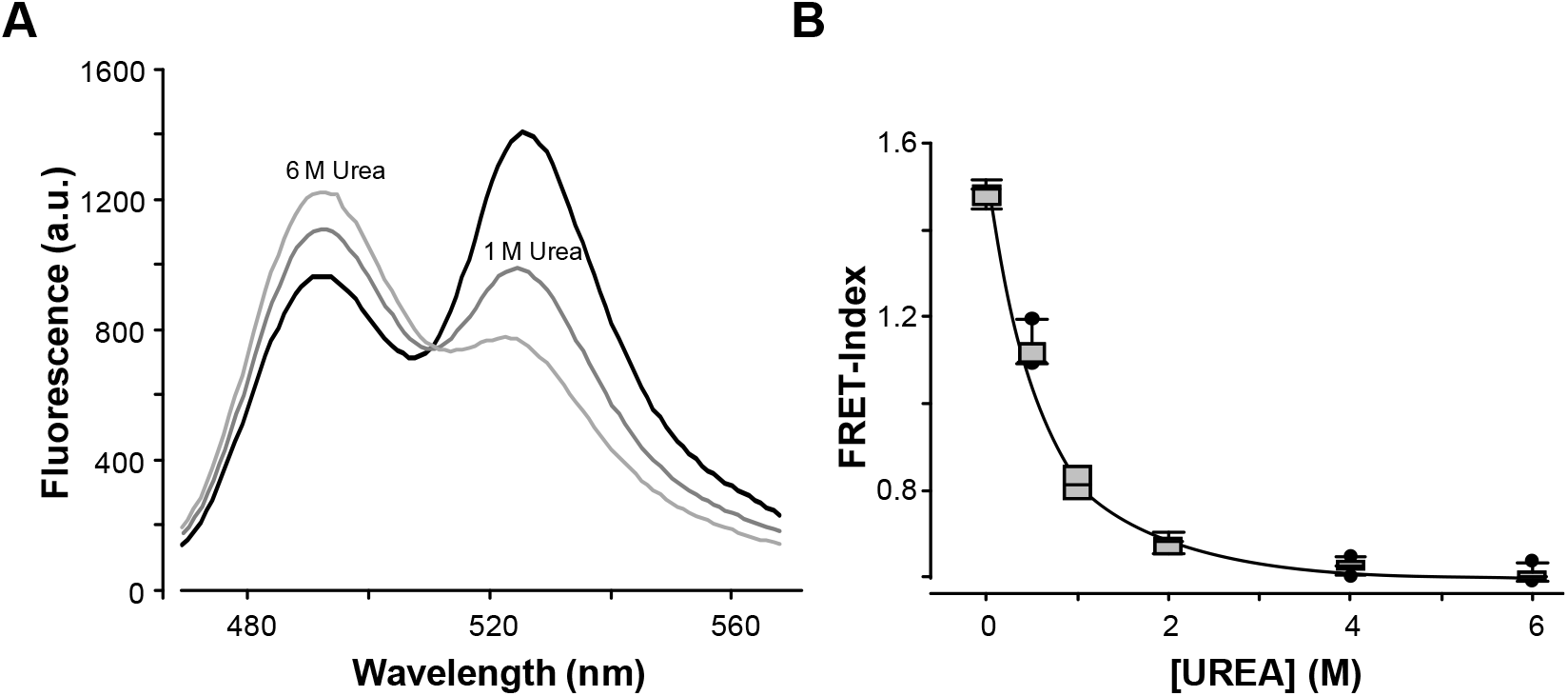
**A.** Emission spectra of the purified mTFP1-AB-mcpVenus/CaM complex in the presence of increasing concentrations of the denaturant urea. **B.** FRET index values expressed as the ratio mcpVenus/mTFP1 peak emission (528/492 nm) ratio from spectra as a function of urea concentration.

**Supplemental Figure 5, related to Figure 5.**
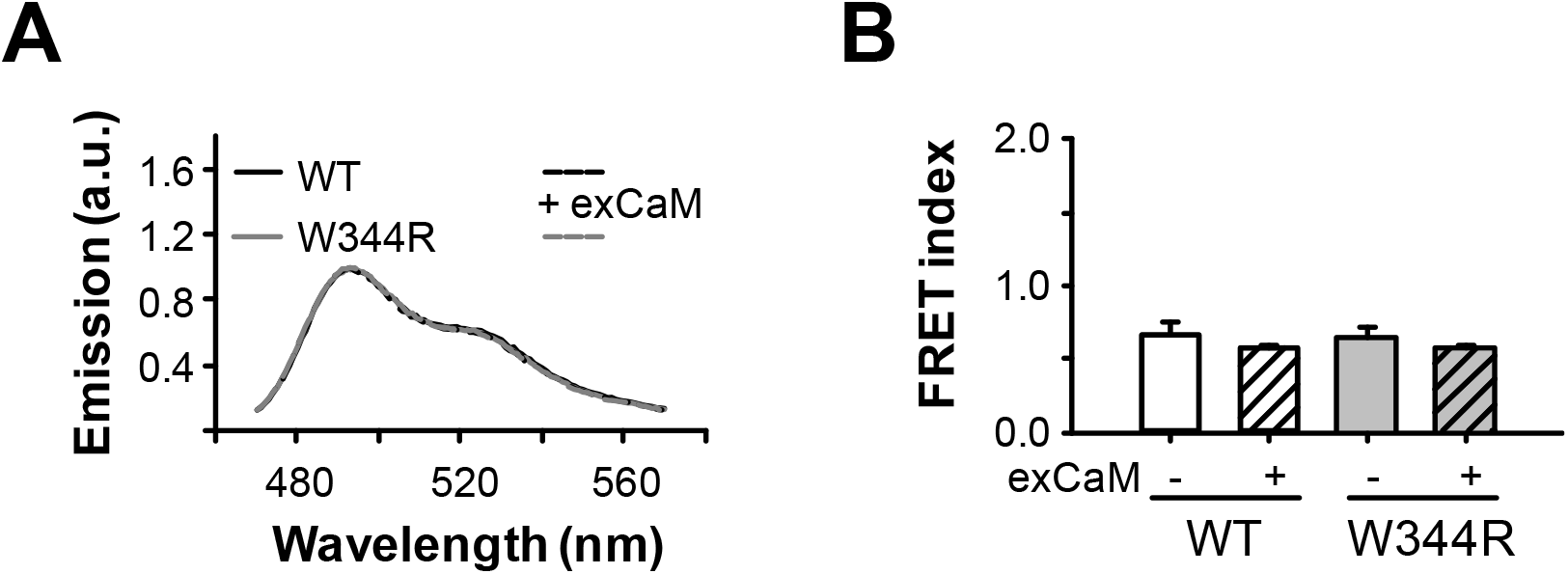
**A.** Emission spectra of the soluble WT (black lines) and W344R (grey lines) proteins at ~6 μM translated in CaM-free non-denaturing conditions. An excess (100 μM) exogenous-CaM (exCaM) was added to each sample (dotted lines) and the emission spectra were measured after 24 hours. **B.** FRET index values expressed as the ratio mcpVenus/mTFP1 peak emission (528/492 nm) from spectra as in A.

**Supplemental Figure 6, related to Figure 5.**
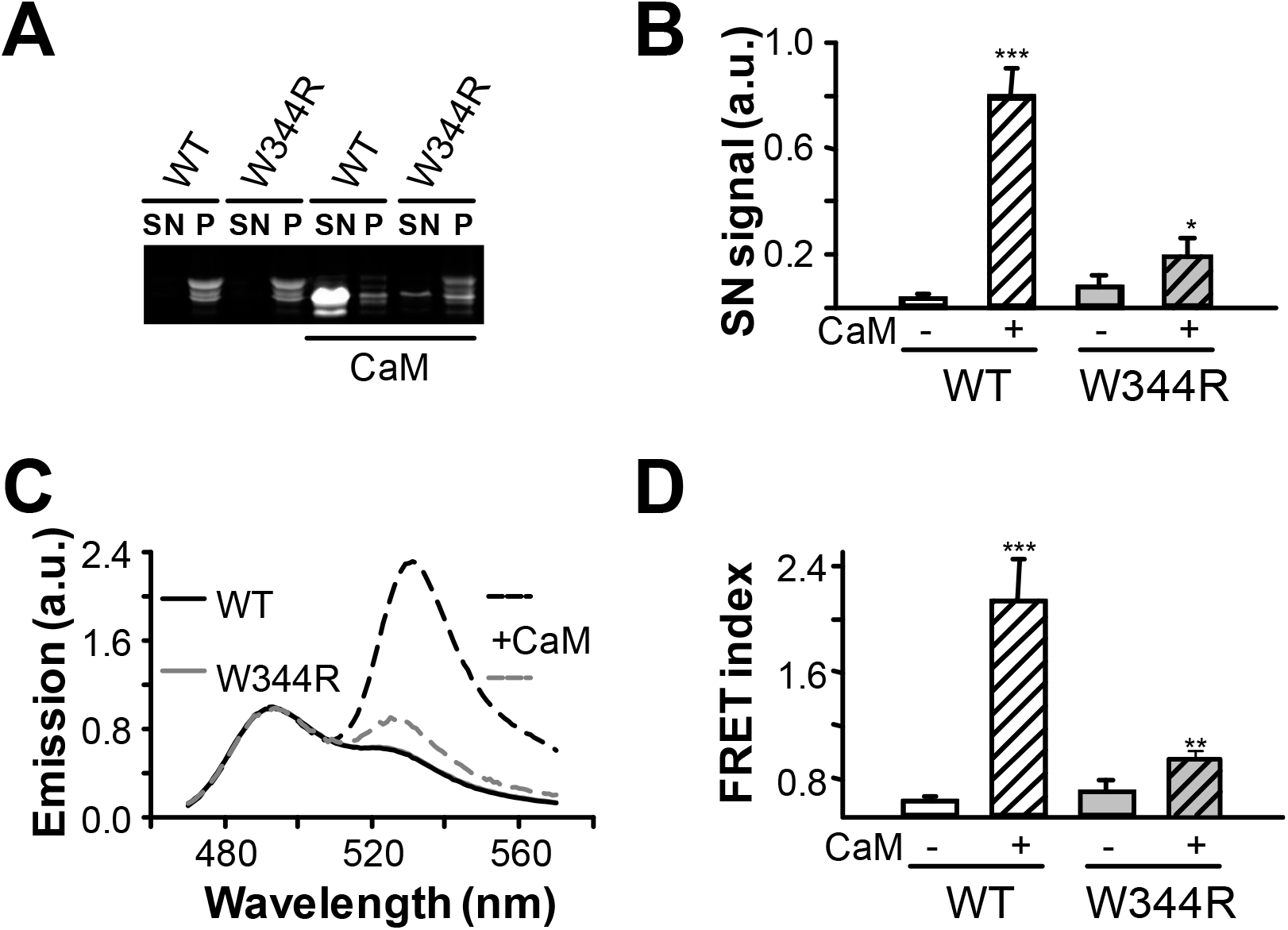
**A.** Fluorescent image of a pseudo-native SDS-PAGE of bacterial extracts of cells expressing WT or W344R biosensors, expressed at 18°C. Proteins were co-expressed (right columns) or not (left columns) with CaM. Soluble (supernatant; SN) and insoluble (pellet; P) protein fractions were separated, and loaded as indicated. **B.** Fluorescence intensity of the supernatant band of SDS-PAGE gel (n = 3). **C.** Emission spectra of the soluble fraction of WT (black lines) and W344R (grey lines) proteins expressed alone (solid lines) or co-expressed with CaM (dashed lines). **D.** FRET index values expressed as the ratio YFP/CFP peak emission (528/492 nm) ratio from spectra as in C.

**Supplemental Figure 7, related to Figure 5.**
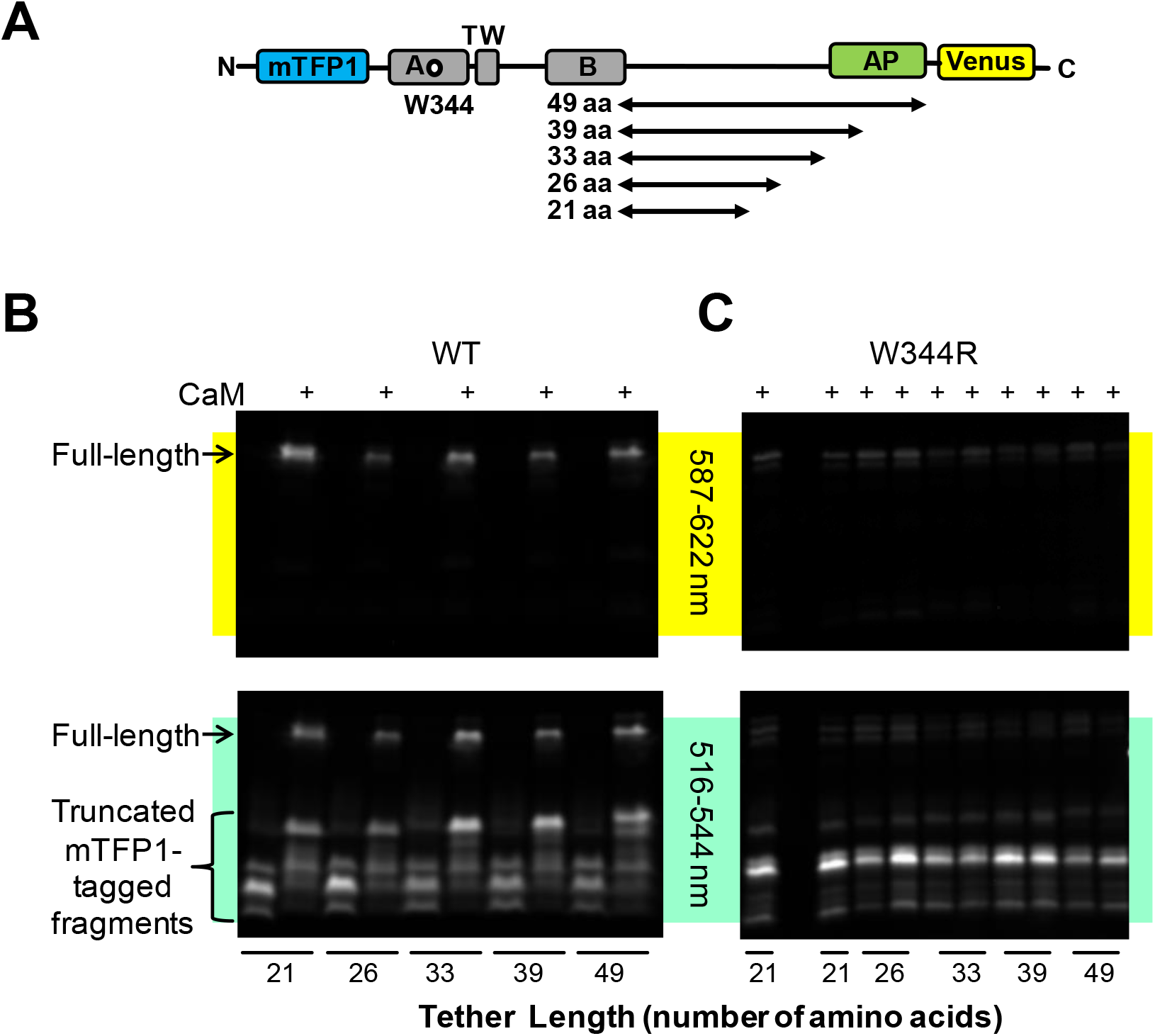
**A.** Schematic representation of the constructs used for *in vivo* translation. The CRD was cloned upstream of the SecM arresting peptide (AP) sequence with tethers of increasing length, ranging from 21 to 49 amino acids from the C-terminal conserved Pro of the SecM AP where translational stalling takes place. **B.** Fluorescent images of a pseudo-native SDS-PAGE of bacterial extracts expressing WT-AP construct with and without CaM, with tether lengths indicated at the bottom. A 605BP35 filter was used to isolate emission from mcpVenus on the image at the top, whereas the image at the bottom, a 530BP28 filter was used to detect emission from both mTFP1 and mcpVenus. **C.** Fluorescent images as in B of bacterial extracts expressing W344R-AP constructs in the presence of CaM. The second line was not loaded.

**Supplemental Figure 8, related to Figure 6.**
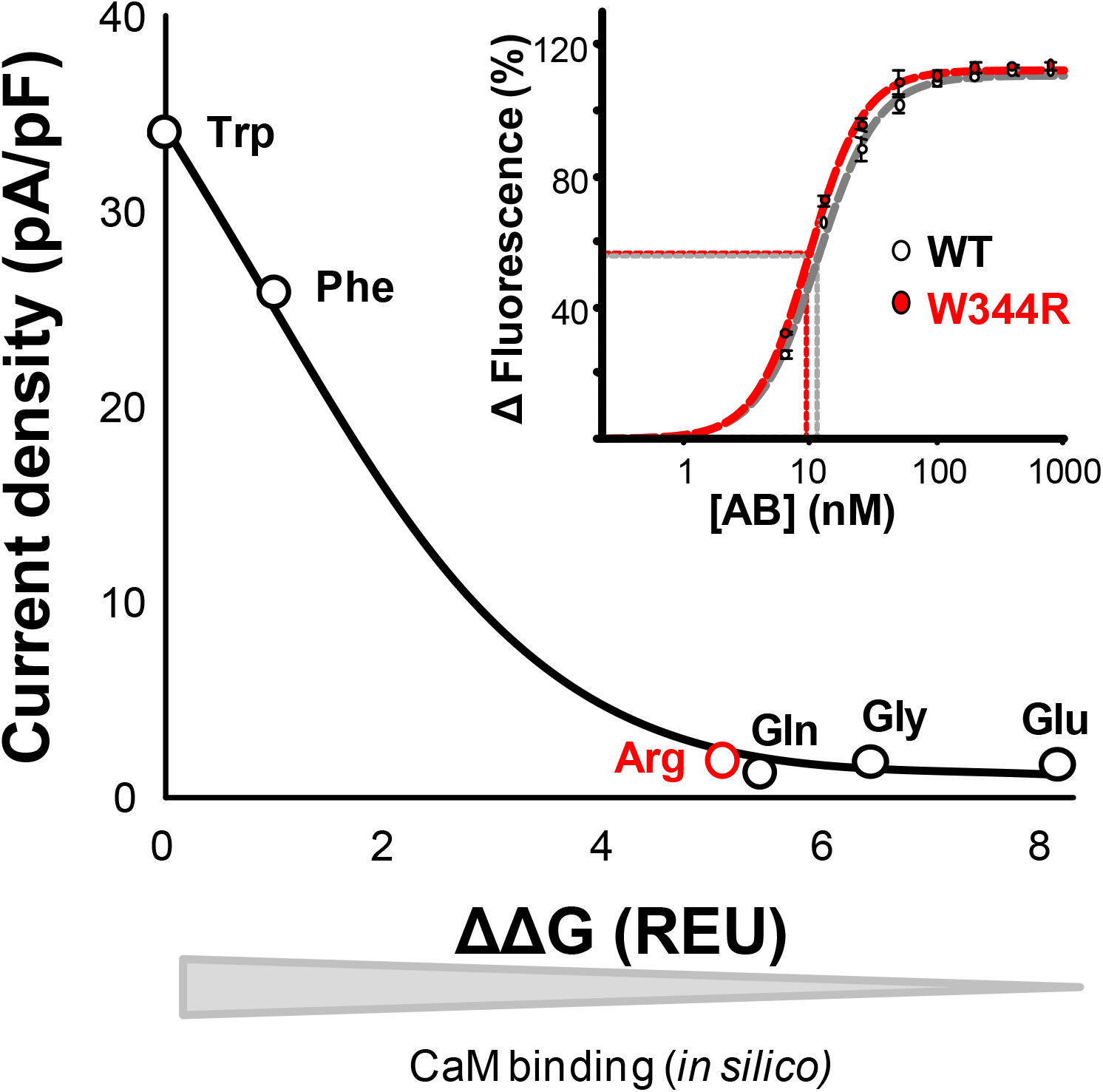
Relationship between current densities of homomeric K_v_7.2 channels carrying the indicated mutations at position 344 and the computed binding energies in Rosetta Energy Units (REU). It is expected that the higher the value, the weaker the predicted affinity for CaM. However, the measured apparent binding affinity is 13% more favorable for the W344R than WT (Inset, from [1]).

**Supplemental Figure 9, related to Figure 6.**
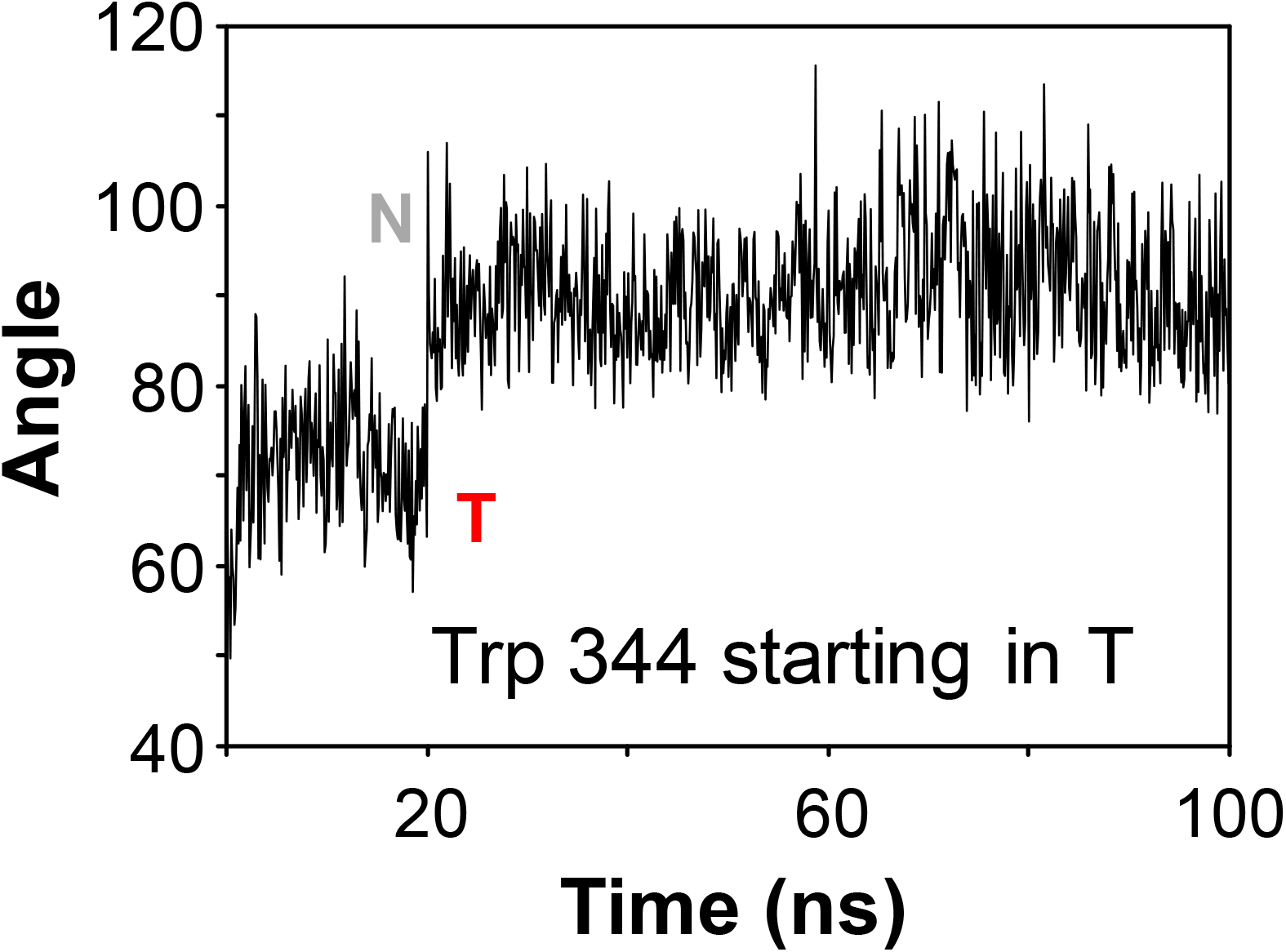
Time series of the angle of Tryptophan 344 through a molecular dynamics simulation of the K_v_7.2 WT CRD forced to start in T configuration. Note that for the first 20 ns, tryptophan 344 remains in T and then exhibits a conformational change towards the more stable N configuration.

